# Alterations in type 2 dopamine receptors across neuropsychiatric conditions: A large-scale PET cohort

**DOI:** 10.1101/2023.06.05.543660

**Authors:** T. Malén, S. Santavirta, S. De Maeyer, J. Tuisku, V. Kaasinen, T. Kankare, J. Isojärvi, J. Rinne, J. Hietala, P. Nuutila, L. Nummenmaa

**Affiliations:** Turku PET Centre, University of Turku, Turku, Finland; Turku PET Centre, Turku University Hospital, Turku, Finland. Electronic address; Turku PET Centre, University of Turku, Finland; Turku PET Centre, Turku University Hospital, Turku, Finland; University of Antwerp, Antwerp, Belgium; Turku PET Centre, Turku University Hospital, Turku, Finland; Clinical Neurosciences, University of Turku, Turku, Finland; Neurocenter, Turku University Hospital, Turku, Finland; Turku PET Centre, University of Turku, Turku, Finland; Turku PET Centre, Turku University Hospital, Turku, Finland; Department of Psychiatry, University of Turku and Turku University Hospital, Turku, Finland; Turku PET Centre, University of Turku, Turku, Finland; Department of Endocrinology, Turku University Hospital, Turku, Finland; Turku PET Centre, University of Turku, Turku, Finland; Turku PET Centre, Turku University Hospital, Turku, Finland; Department of Psychology, University of Turku, Turku, Finland

**Keywords:** dopamine type 2 receptor, neuropsychiatric conditions, positron emission tomography, Bayesian mixed effects modeling

## Abstract

**PURPOSE:** Aberrant dopaminergic function is linked with motor, psychotic, and affective symptoms, but studies have typically compared a single patient group with healthy controls. METHODS: Here, we investigated the variation in striatal (caudate nucleus, nucleus accumbens, and putamen) and thalamic type 2 dopamine receptor (D_2_R) availability using [^11^C]raclopride positron emission tomography (PET) data from a large sample of 437 humans including healthy controls, and subjects with Parkinson’s disease (PD), antipsychotic-naïve schizophrenia, severe violent behavior, pathological gambling, depression, and overweight. We analyzed regional group differences in D_2_R availability. We also analyzed the interregional correlation in D_2_R availability within each group. RESULTS: Subjects with PD showed the clearest decline in D_2_R availability. Overall, the groups showed high interregional correlation in D_2_R availability, while this pattern was weaker in violent offenders. Subjects with schizophrenia, pathological gambling, depression, or overweight did not show clear changes in either the regional receptor availability or the interregional correlation. CONCLUSION: We conclude that the dopaminergic changes in neuropsychiatric conditions might not only affect the overall receptor availability but also the connectivity between the regions. The region-specific receptor availability more profoundly links to the motor symptoms, while the between-region connectivity might be disrupted in violence.

**Highlights:** We compared human striatal and thalamic type 2 dopamine receptor (D_2_R) availability between healthy controls, and subjects with Parkinson’s disease (PD), antipsychotic-naïve schizophrenia, severe violent behavior, pathological gambling, depression, and overweight.

We present the mean brain maps of group specific D_2_R availabilities in NeuroVault (https://neurovault.org; https://identifiers.org/neurovault.collection:12799).

Dopamine type 2 receptor availability is lowered in PD in caudate nucleus, nucleus accumbens and thalamus.

Subjects with severe violent behavior had decreased correlation between the striatal and thalamic D_2_R availability.

Altered regional D_2_R availability in the striatum and thalamus is linked with motor disorders, while lowered interregional connectivity in D_2_R might relate to violence.

**KEY POINTS:** QUESTION: Are there differences in the striatal (caudate nucleus, nucleus accumbens, and putamen), and thalamic D_2_R availability in a sample including healthy controls, and subjects with Parkinson’s disease, antipsychotic-naïve schizophrenia, severe violent behavior, pathological gambling, depression, and overweight?

PERTINENT FINDINGS: Based on this register-based study of a large historical sample (n=437), Parkinson’s disease links to changes in the regional receptor availability, while in severe violent behavior, the correlation between regional receptor availabilities might be lowered. No clear receptor changes were observed in overweight.

IMPLICATIONS FOR PATIENT CARE: Based on our data of striatal and thalamic type 2 dopamine receptors, region-specific changes are linked with motor disorders, while lowered between-region correlation might relate to violence.

## Introduction

Neurotransmitter dopamine is centrally involved in motor, motivational and emotional processes [1]. The four major dopaminergic pathways are i) nigrostriatal pathway running from substantia nigra to the dorsal striatum (i.e. caudate nucleus and putamen), ii) mesolimbic pathway from ventral tegmental area (VTA) to the ventral striatum (i.e. nucleus accumbens), iii) mesocortical pathway from VTA to prefrontal and cingulate cortex, and iv) tuberoinfundibular pathway from hypothalamus to pituitary gland [2, 3]. The nigrostriatal pathway is critically involved in motor function and Parkinson’s disease (PD) [4], while the mesolimbic pathway (also known as the reward pathway) contributes to motivation [2, 5]. Particularly, the striatal type 2 dopamine receptor (D_2_R) availability? has been linked with a variety of neuropsychiatric symptoms [6–8].

Lowered dopamine transmission in the nigrostriatal pathway is associated with motor symptoms (e.g. rigidity, and bradykinesia) in PD [8], while hyperactivity of the dopamine neurons in the mesolimbic track might promote psychotic symptoms in schizophrenia [4, but see also 9]. Antipsychotic-naïve schizophrenia patients with first-episode psychosis show lowered availability of thalamic D_2_Rs [10], possibly also linked to elevated presynaptic dopamine release, observed at least in the closely located striatum [11]. However, while increased presynaptic dopamine synthesis and release is characteristic for schizophrenia, only some patients have altered postsynaptic D_2_R, pointing towards subtypes of schizophrenia [12].

Dopamine also contributes to impulsive behavior, such as aggressive outbursts and drug abuse [13–16]. Mesolimbic dopamine pathway and upregulated striatal dopamine function is critical for aggressive behavior [15]. Hyperactive striatal dopaminergic function is also consistently linked with acute effects of drug abuse in addicted subjects, who overall show diminished dopamine and D_2_Rs in the striatum [16]. Baseline downregulation of dopamine in addictions might associate with their commonly comorbid affective symptoms [17], such as anhedonia (lack of pleasure) and dysphoria (dissatisfaction), that are coherently connected to diminished mesocorticolimbic dopamine [18]. Overall, affective symptoms are involved in numerous neurological and psychiatric conditions, including PD, schizophrenia and depression [18, 19].

Finally, drugs that enhance striatal dopaminergic neurotransmission effectively treat motor symptoms [20] but can also induce psychotic symptoms, impulsivity, and addictive behavior (e.g. gambling and overeating) [4, 5, 21]. Conversely, neuroleptics, pharmacological treatments that inhibit the striatal dopaminergic function by blocking the D_2_Rs, reduce psychotic symptoms [18, 22] and aggressive behavior [15], but induce Parkinsonian-like side effects, such as rigidity and bradykinesia [3, 9, 23, 24], and anhedonia [18]. Particularly the antipsychotics that affect the nigrostriatal track in addition to the mesolimbic track might be the ones with motor side-effects [23], further supporting the centrality of the nigrostriatal track in motor functions.

Taken together, existing data suggest that elevated striatal dopamine activity contributes to psychotic symptoms, impulsivity, aggressive outbursts, and acute effects of reward (namely intense hedonia / pleasure), while lowered dopamine activity relates to disturbed motor functions, and blunted affect (e.g., anhedonia, diminished reward experiences). Because several neuropsychiatric conditions share symptoms [14, 19], perturbations in the dopamine system might link to specific neuropsychiatric symptoms regardless of the diagnosis. To resolve the links between the pathologies with shared dopamine-regulated symptoms of voluntary movement, motivation, and affect, it is crucial to study them together. Only this allows unravelling whether differential patterns of dopaminergic dysfunctions underlie different neuropsychiatric conditions. Here, by neuropsychiatric we refer to a combined set of pathologies with an organic (neurological) and/or psychological basis.

We investigated the differences in the striatal (caudate nucleus, nucleus accumbens, and putamen), and thalamic D_2_R availability in a large (n= 437) historical sample of subjects using positron emission tomography (PET) data with radioligand [^11^C]raclopride acquired in resting baseline state. The dopaminergic function includes tonic and phasic dopamine firing [25]. Because our data pertain to baseline, resting state scans, these data reflect predominantly tonic rather than phasic dopaminergic function. Our sample included healthy controls, and subjects with PD, antipsychotic-naïve schizophrenia, severe violent behavior (violence), pathological gambling (gambling), depression, and overweight. We also analyzed between-region “connectivity” (correlation) of the D_2_R availability in each group and assessed the effects of age and sex on D_2_R availability in i) the healthy subjects and ii) the other six groups (i.e. groups of interest) together.

## Material and Methods

### Data

Basic subject information is shown in **Table 1**. This retrospective register-based study includes 437 subjects (294 males, 143 females), including healthy controls (n= 239), and subjects with PD (n= 60), schizophrenia (n= 7), severe violent behavior, i.e. prison sentence for violence (n= 10), pathological gambling (n= 12), depression (n= 12), and obesity/overweight (n= 97) who had undergone [^11^C]raclopride PET scan. The age range of the subjects was 19-82 years. The data were retrieved from the Turku PET Centre Aivo database (https://aivo.utu.fi), including data of imaging studies conducted at the site. See supplementary material for **Original publications whose data are used in the current study**. The images were acquired between the years 1989 and 2019 with six different PET scanners (GE Advance, HR+, HRRT, GE Discovery VCT PET/CT, GE Discovery 690 PET/CT, Ecat 931). The scanners are described in detail in our previous work [26], and the distribution of the different PET scanners across the subject groups are given in supplementary **Table S1**.

**Table 1.** Age of the sample groups separately for males and females. This is the final sample of 437 subjects (excluded subjects are not reported here).

|  | Males (n= 294) |  |  |  | Females (n= 143) |  |  |  |
| --- | --- | --- | --- | --- | --- | --- | --- | --- |
|  | N | Mean | SD | Range | N | Mean | SD | Range |
| Healthy controls | 174 | 28 | 11 | 19-78 | 65 | 41 | 16 | 19-82 |
| Parkinson's disease | 31 | 62 | 12 | 34-82 | 29 | 60 | 11 | 41-81 |
| Schizophrenia | 3 | 22 | 3 | 19-25 | 4 | 36 | 13 | 24-53 |
| Violence | 10 | 31 | 7 | 24-46 | 0 | - | - | - |
| Pathological gambling | 12 | 32 | 8 | 23-49 | 0 | - | - | - |
| Depression | 3 | 38 | 10 | 25-48 | 9 | 41 | 5 | 34-50 |
| Overweight | 61 | 29 | 9 | 19-56 | 36 | 43 | 13 | 23-74 |

We retrieved the following subject information at the time of each PET scan: subject age, sex, body mass index (BMI, available for 67% of the subjects) and group, as well as the injected dose and the PET scanner used to acquire the imaging data. Although we do not have detailed medication information for all the subjects, the exclusion criteria for the original research projects where that the subjects were drawn from contained medication affecting the central nervous system. The schizophrenia group was antipsychotic-naïve, and the depression group drug-free (mainly drug-naïve subjects with moderate depressive episode). The PD sample contained both unmedicated and medicated patients and given the mean disease duration of 10 years (median= 9 years, range from 0 to 29 years, see supplementary **Figure S1** under **D_2_R availability through PD duration**), most of the patients were probably medicated at the time of imaging. The scans were, however, performed after a short washout period (subjects were withdrawn from medication at least from the night before the scan). We set the BMI cutoff between healthy control and overweight groups to 25 [27], and the BMI range of the remaining subjects (including subjects with PD, schizophrenia, severe violent behavior, pathological gambling, and depression) was 19-49 (regards 67% of the total sample for who we had BMI information available, see supplementary **Table S2**). More detailed symptoms, alcohol use, or smoking status were not used in the analysis, as this information was not systematically available for all the subjects.

Three subjects with low injected dose < 90 megabecquerel (MBq) and seven without dose information were excluded from the sample to avoid low signal to noise ratio. The excluded subjects were from healthy control, PD, schizophrenia, and overweight groups. Due to lost or noisy data from interrupted PET scanning, we discarded eight subjects (including subjects from PD, schizophrenia, and severe violent behavior groups) whose kinetic model fits were inadequate. Altogether 18 subjects were excluded from the data due to aforementioned reasons and the final sample size was 437.

### PET data preprocessing and kinetic modelling

PET image preprocessing and kinetic modelling were carried out in MATLAB [28] using an in-house brain image processing pipeline Magia [29] (https://github.com/tkkarjal/magia). We spatially normalized the dynamic PET images into MNI152 space with an in-house [^11^C]raclopride PET-template created using a subset (n=187) of healthy control subjects, because only a subset of the sample had magnetic resonance (MR) image available (n=249) [29] (see **Validation of PET atlas based spatial normalization of the [11C]raclopride binding estimates** and **Figures S2-S3** in supplementary material). Four bilateral regions of interest (ROI) from Harvard-Oxford atlas, caudate nucleus (caudate), nucleus accumbens (accumbens), putamen, and thalamus, were applied to the normalized PET-images. [^11^C]raclopride binding to D_2_Rs in these regions is reliable, while negligible elsewhere [30, 31].

Regional specific binding of [^11^C]raclopride was quantified as nondisplaceable binding potential (BP_ND_) using simplified reference tissue model (SRTM) and cerebellum as a reference region [32]. BP_ND_ resembles receptor availability, being a measure of receptor density, affinity, and the receptor occupancy by endogenous dopamine [33, 34]. Parametric BP_ND_ images were also calculated for illustrative purposes with basis function implementation of SRTM (bfSRTM) with 300 basis functions. Lower and upper bounds for theta parameter were set to 0.082 1/min and 0.6 1/min. Before the parametric image calculation, the dynamic PET images were smoothed using Gaussian kernel with 4 mm full width at half maximum to reduce the effect of noise in the voxel-level bfSRTM fit. In the analyses, we used the mean of left and right hemisphere BP_ND_ as the regional estimate for each subject. Data on hemispheric symmetry in BP_ND_ is presented in supplementary **Table S3** and **Figure S4**.

### Statistical analysis

Statistical analysis was carried out in R [35]. We assessed the normality of the data with Shapiro-Wilk test with package stats [35]. We analyzed the regional data using Bayesian modeling tools of brms [36–38] utilizing rstan [39]. We analyzed the interregional BP_ND_ correlation in each subject group using correlation matrix with corrplot [40], and Mantel test [41] for similarity of two matrices using ape [42]. We used R [35], particularly ggplot2 [43], ggsci [44], bayesplot [45], superheat [46], corrplot [40], and GGally [47], MATLAB [48], and MricroGL (https://www.nitrc.org/projects/mricrogl) to visualize the data and findings.

#### Regional group differences in D_2_R availability

We used linear mixed effects regression to compare regional D_2_R availability across the seven subject groups. In the analysis, we adjusted for the scanner used in the image acquisition, and the subject age and sex that affected striatal BP_ND_ estimates in healthy controls [26].

The regional binding potentials (BP_ND_) were modeled as a dependent variable. We analyzed the group differences in BP_ND_ in caudate, accumbens, putamen, and thalamus, while adjusting for age, sex, and scanner. Specifically, we applied group, standardized age, and sex as fixed effects and allowed these effects to vary between ROIs (regionally varying a.k.a. random slopes). We allowed the BP_ND_ intercept to vary between subjects, scanners, ROIs, and the scanners within each ROI (random intercepts for subjects, scanners, ROIs, and scanner-ROI combinations). For the residual variances, we similarly applied random intercepts for scanners, ROIs, and the scanner-ROI combinations, but not for subjects as we did not expect the residual variances to depend on subject. We validated the model by comparing it with three other candidate models (see **Model diagnostics and comparison** in supplementary material).

The age effect was calculated for the whole sample together (and not separately for each subject group by interaction of age and group), because we wanted to maximize the age range and the sample size in the estimation of the age effect, while some of the studied groups had relatively limited age range (e.g. schizophrenia 19-25 years). Group-specific age effects were also not hypothesized as e.g. prior studies suggest that neural changes in PD do not reflect ‘accelerated aging’ [49]. Similarly, the sex effect was calculated for the whole sample because some of the studied groups consisted solely of males. Nevertheless, of all the studied groups, the healthy controls hold the greatest number of observations altogether and separately for males and females, as well as the widest age range. Thus, the age and sex effects observed in the healthy control group will weight a relatively lot in the regression analysis due to their strong representation in relation to the other groups.

Shapiro Wilk testing [35] did not clearly support the normality of region-specific BP_ND_ in the original nor the logarithmic scale. We decided to use the log-transformed instead of the original-scale BP_ND_ as a dependent variable in the regression model, because log-transformed dependent variable allows us to directly estimate the relative (e.g. 10%) rather than absolute (e.g. 1 unit) change in BP_ND_ between predictor levels, particularly the group specific intercepts. Relative change is more intuitive, while BP_ND_ is a ratio that describes receptor availability, and is not a direct measure of absolute receptor density [50].

We used linear regression, because based on our previous assessment [26], the relationship between age and logarithmic BP_ND_ is well approximated by a linear function in a healthy sample, that is the majority of the current sample, and as stated previously, we are not aware of group-specific age-effects. Further, modeling higher-degree associations increases the risk of overfitting, potentially resulting in poor generalization of the results [51]. For the remaining fixed predictors (group and sex), the estimated parameters are intercepts and not slopes.

In the absence of appropriate priors for Bayesian statistical inference from independent data, we gave a normal distribution prior (with expected value= 0 and variance= 1) for the fixed effects (age, sex, group). Such zero-centered normal prior assigns symmetrical probabilities for both positive and negative regression coefficients and does not restrict the parameter to certain values (i.e., no regression coefficient is given zero probability), still regularizing the model’s fit. For the standard deviations of the random effects, we set the same normal prior. As standard deviation is always positive, brms-package automatically restricts this prior to positive values, making it a half-normal distribution [36–38]. For the rest of the modeling parameters, we used weakly informative default priors of brms [36]. The model was carried out using Markov Chain Monte Carlo sampling with each four chains running 4000 iterations (including 1000 warm-up samples).

#### Validation of the age and sex effects on regional D_2_R availability

For validation purposes, we assessed the age and sex effects in the current sample. We tested the replicability of the age and sex effects [26] in the i) healthy control sample (almost identical sample to our earlier work [26]) and ii) groups of interest together (subjects with PD, schizophrenia, severe violent behavior, pathological gambling, depression, and overweight) (**Validation of the age and sex effects on regional D_2_R availability** in supplementary material).

#### Between-region correlation of the D_2_R availability

We computed a 4×4 between-region BP_ND_ correlation matrix for each subject group. These matrices were computed with the original binding estimates, because the data (group and region specific BP_ND_) were not normally distributed (Shapiro-Wilk), and the rank-order based Spearman correlation readily accounts for monotonic associations in the data and is unaffected by log transformations. Using Spearman and not Pearson correlation is also supported by the limited sample sizes in some of the studied groups. One-tailed testing was used as regional coupling estimates for D_2_R are positive rather than negative [26].

We tested for the similarity of each correlation matrix pair (e.g. correlation matrix of healthy controls compared to correlation matrix of PD patients) using one-tailed Mantel tests with default number (999) of permutations [42]. These analyses allowed us to assess, whether interregional D_2_R coupling is altered in neuropsychopathology, and whether different neuropsychiatric conditions share or differ in their striatal and thalamic ‘connectivity’ patterns.

## Results

### Descriptives

Group-specific D_2_R mean, and coefficient of variance maps are shown in **Figure 1**. Based on the **Figure 1**, the groups show broadly similar receptor distribution in the subject groups, and the binding level shows greatest variation in the PD group. Across groups, the coefficient of variation appears greatest in thalamus. The mean D_2_R maps are also available in NeuroVault (https://identifiers.org/neurovault.collection:12799). Mean regional non-transformed BP_ND_ estimates for each group are shown in **Figure 2**. D_2_R availability decreases through PD duration, please see **D_2_R availability through PD duration** and **Figure S1** in the supplementary material.

**Figure 1.**
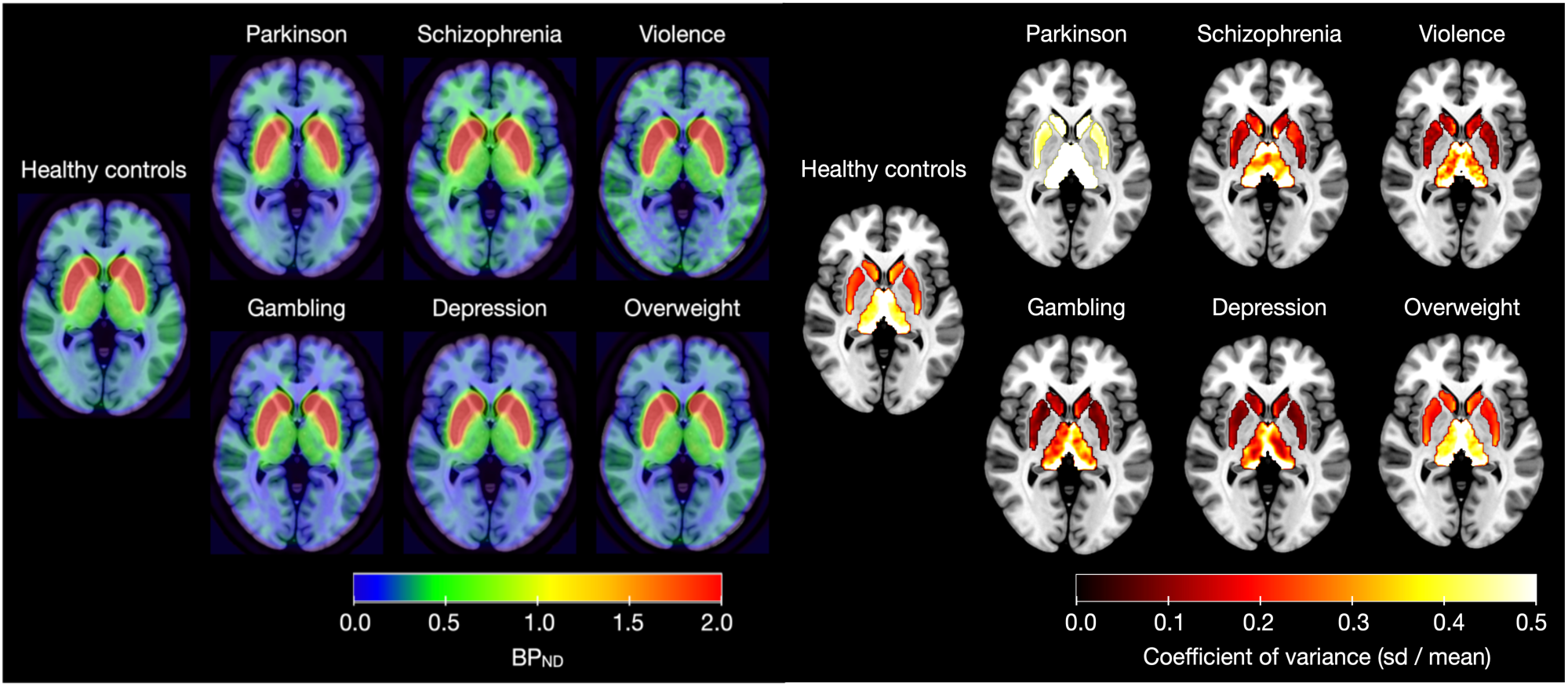
Assessment of the original-scale [^11^C]raclopride BP_ND_ estimates in the subject groups. Left: Mean binding. Right: Binding coefficients of variation (standard deviation / mean) within the regions of interest (Harvard-Oxford atlas masks) on an MNI-template.

**Figure 2.**
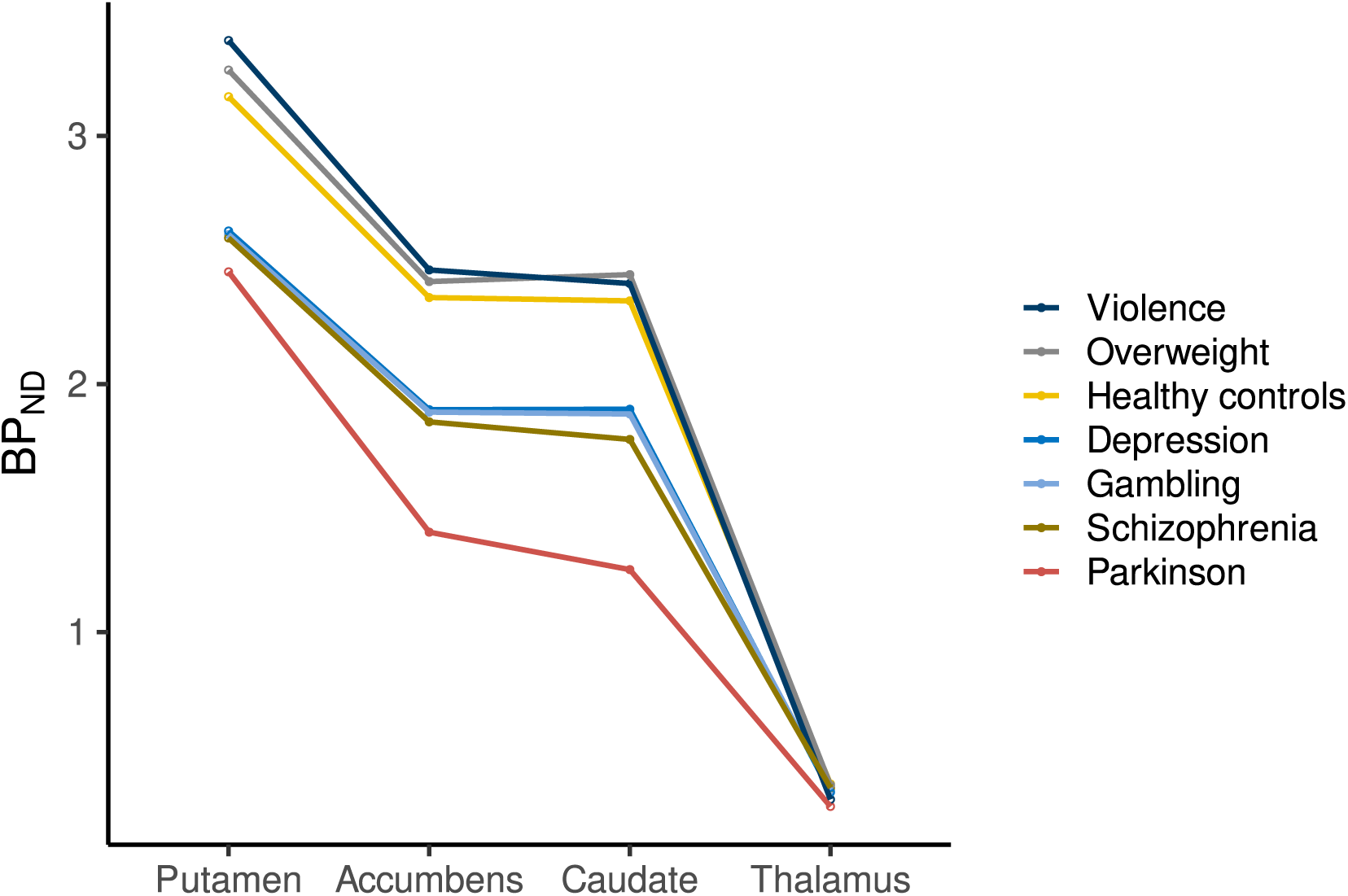
The group specific original-scale BP_ND_ profiles. The lines connect the group specific mean points to aid in visualization of the group-specific binding profiles.

### Regional group differences in D_2_R availability

Bayesian linear mixed effects modeling (**Figure 3**), adjusting for age, sex, and scanner, revealed that compared to the other groups, PD patients had approximately 10-20% lower [^11^C]raclopride BP_ND_ (values transformed back to original scale) in caudate, accumbens and thalamus, while the BP_ND_ in putamen was approximately 10% higher (**Figure 4**). The other groups of interest (schizophrenia, severe violent behavior, pathological gambling, depression, and overweight) did not show clear differences in regional D_2_R availability when compared with healthy controls or each other. However, subjects with schizophrenia and severe violent behavior showed more support for lowered than elevated D_2_R availability (more probability mass below zero), although the 80% posterior intervals (i.e. probability mass) crossed zero. Pathological gambling, depression and overweight groups showed the least support for altered D_2_R availability with practically non-existent difference compared to healthy controls.

**Figure 3.**
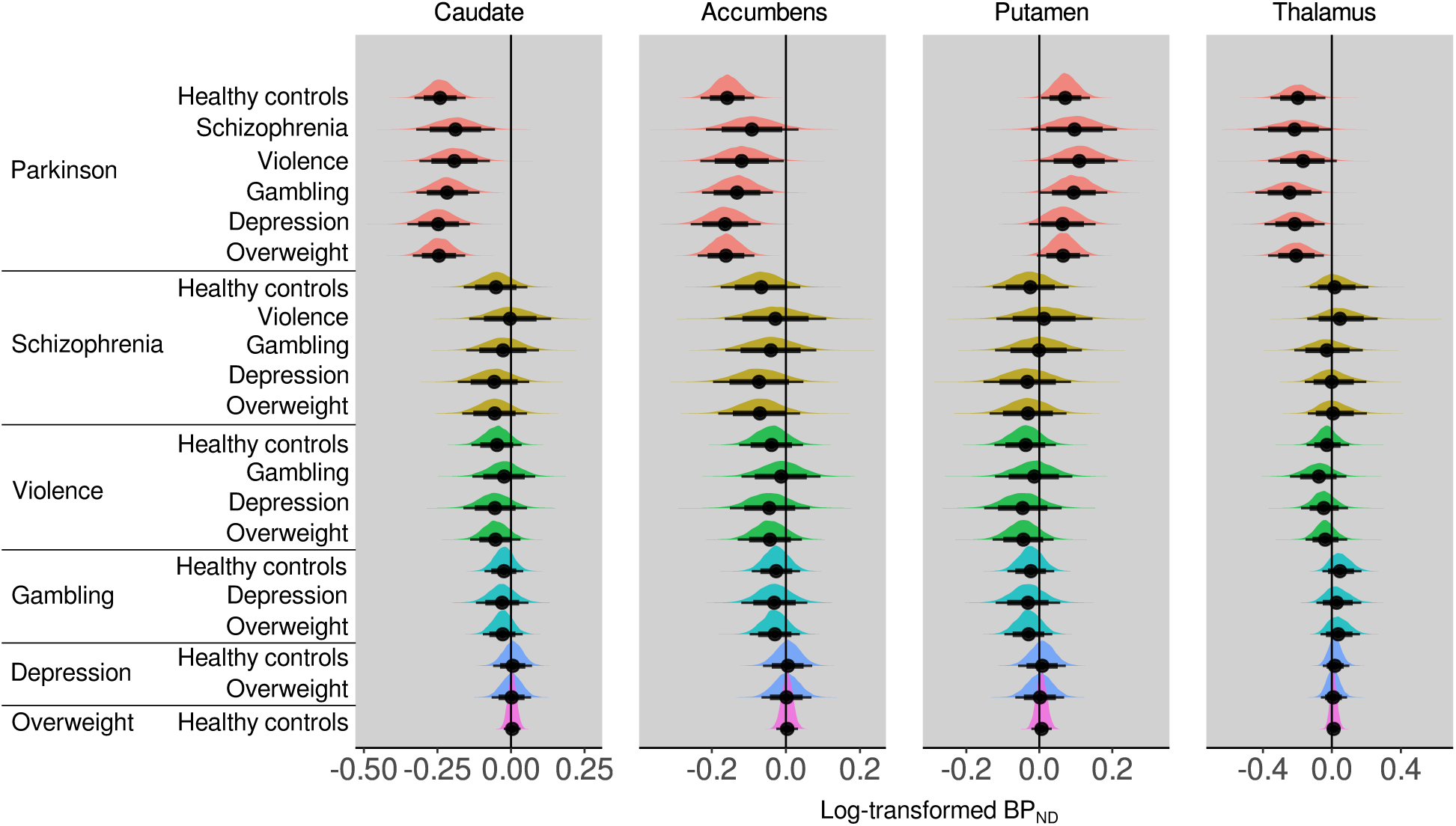
The figure shows colored posterior probability distributions (posteriors), as well as their median (point), 80% (thick line) and 95% (thin line) posterior intervals, describing the between-group differences in log-transformed BP_ND_. Posterior located on the negative side of the zero line suggests lower log-BP_ND_ in the reference group on the left compared with the group on the right.

**Figure 4.**
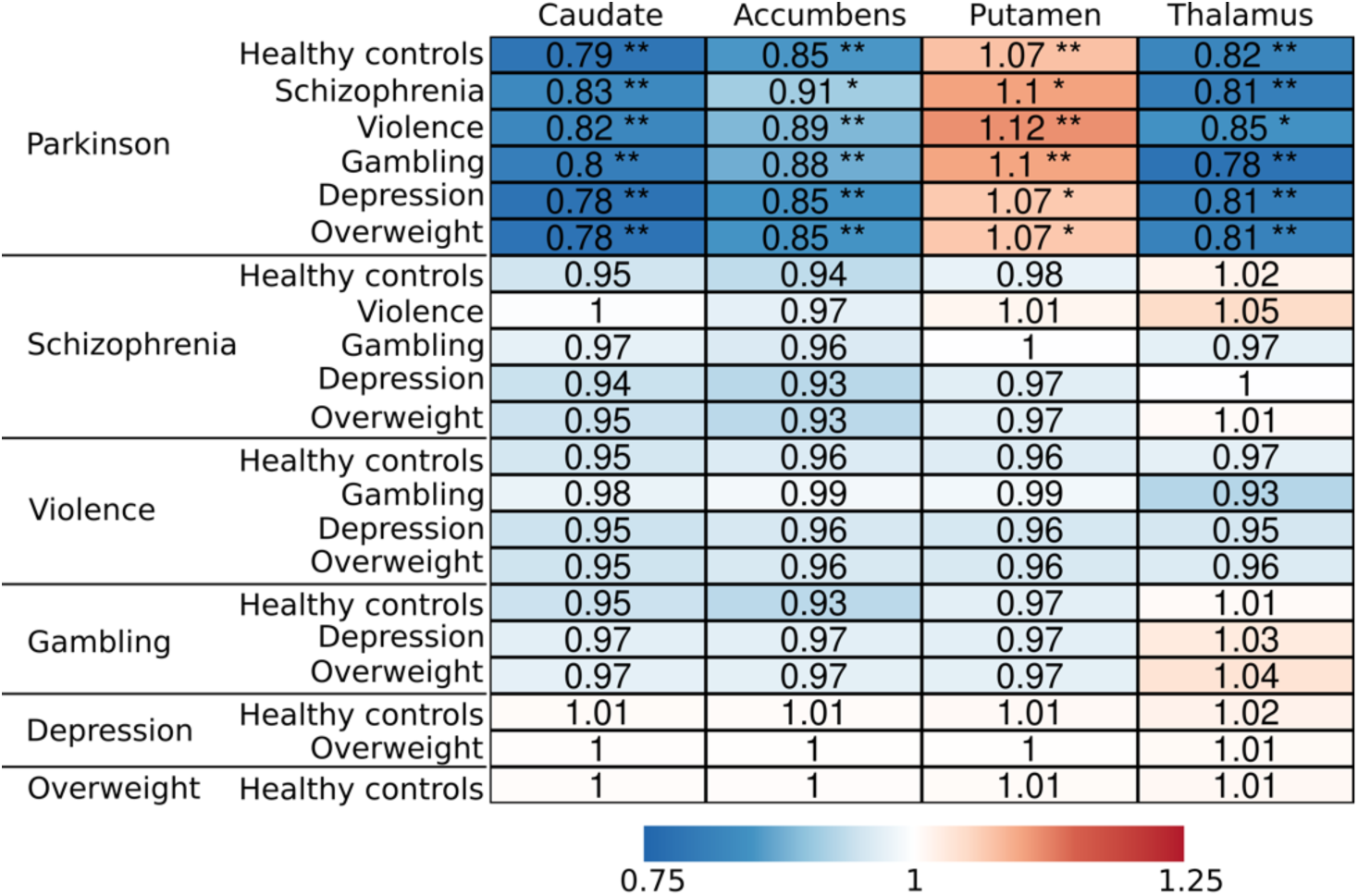
Summary of the regional group comparisons. The values represent the (median) proportional BP_ND_ (values transformed back to original scale) of the reference group (on the left) compared with the group on the right. Asterisks represent comparisons were 80% (*) and 95% (**) posterior interval did not cross zero.

### Validation of the age and sex effects on regional D_2_R availability

Similarly as in our previous analysis with healthy subjects [26], D_2_R availability decreased through age, both in the healthy control group that is almost identical to the sample in our previous work [26] and in the groups of interest. The lower striatal D_2_R availability in males versus females was only found in healthy controls. However, in thalamus, females had higher thalamic availability in both the healthy controls and the groups of interest. The results are illustrated in **Figure S5**. For comparison, **Figure S5** also shows the age and sex effects of the main model and confirms that these effects are roughly similar between the models.

### Between-region correlation of the D_2_R availability

The interregional correlations of original-scale BP_ND_ are shown in **Figure 5**. Additional between-region scatter, and density plots are given in supplementary **Figure S6**. Despite the regional BP_ND_ changes, PD patients showed high between-region BP_ND_ correlation, comparably with healthy controls. Across all groups, the weakest correlations were observed between thalamus and other regions. Subjects with severe violent behavior showed the weakest correlation between the regions.

**Figure 5.**
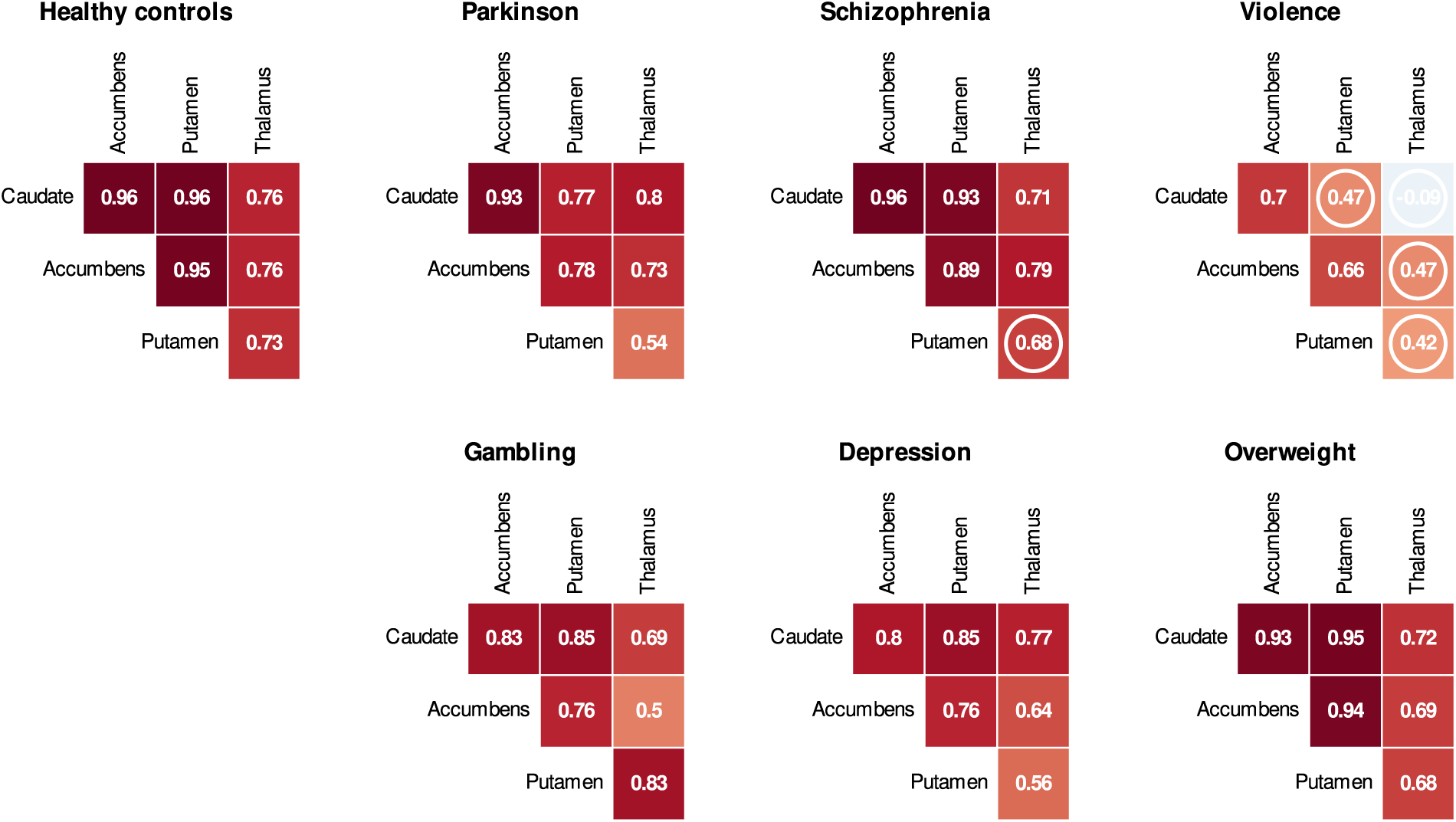
Group specific interregional original-scale BP_ND_ correlations (one-tailed Spearman, p-values > 0.05 marked with a circle, while the higher the p-value, the less support for non-zero correlation).

When we compared the correlation matrixes of the groups (**Figure 5**) with Mantel tests, we observed similarity in the connectivity patterns of the groups. Healthy controls showed some support for similarity with PD (p = 0.04), schizophrenia (p= 0.055), depression (p= 0.037), and overweight (p= 0.04). Further, similarity was suggested between PD and schizophrenia (p= 0.07), PD and depression (p= 0.038), PD and overweight (p= 0.089), and depression and overweight (p= 0.034). The remaining group comparisons, mainly including subjects with severe violent behavior and pathological gambling, did not support similarity, while the p-value was greater than 0.1 in the similarity tests.

Altogether, the groups showed strong dopaminergic connectivity patterns, particularly between the striatal regions. The weakest correlations were most notably manifested as hypoconnectivity of thalamus, where the estimates might be less reliable than in the striatum [31]. The thalamic hypoconnectivity was emphasized in the subjects with severe violent behavior, who had lowest correlation pattern also within the striatum.

## Discussion

Striatal dopamine neurotransmission has a central role in motor functions that are disturbed particularly in PD [52, 53], and motivational processes that are impaired in many neuropsychiatric disorders [13, 15, 16, 18, 19]. Our findings show that in PD, striatal (caudate and accumbens) and thalamic D_2_R availability is downregulated, while interregional correlation might be lowered in severe violent behavior. These results suggest that in the striatothalamic dopamine circuit, overall changes in receptor levels and specific regional decoupling patterns associate to different neuropsychiatric conditions.

### Regional D_2_R changes in Parkinson’s disease

Striatal (caudate and accumbens) and thalamic downregulation in D_2_R availability was most salient in PD patients. There was limited evidence for striatal downregulation also in schizophrenia and severe violent behavior, but even the 80% posterior intervals for these groups overlapped with zero. Our findings support a previous meta-analysis describing that the effect of the striatal D_2_R in schizophrenia is negligible (and possibly linked to the antipsychotic treatment) compared to the presynaptic dopamine synthesis and release [11]. The other groups did not show clear alterations in striatum nor thalamus.

Compared with the other groups, PD patients exhibited both lowered (accumbens, caudate, thalamus) and elevated (putamen) striatal D_2_R availability when adjusting for age and sex. PD subjects had lowered D_2_R availability in nuclei caudate and accumbens, and thalamus. In putamen PD patients had high D_2_R availability, although this effect is not discernible in the raw data where age and sex are not adjusted for.

Although age and group were both accounted for in the regression modeling, downregulation of D_2_R in PD might have been partly misinterpreted as an age effect, as PD patients are on average older than the other subjects (**Table 1**). This possible overcorrection of the age effect might have contributed to the sign switch of the observed alteration of D_2_R in the PD group (positive in the putamen, negative in all other ROIs). However, the age and sex effects were roughly similar in the main model (with all groups included) as in the model of only healthy controls (**Figure S5**). These findings suggest that the low D_2_R availability of PD patients did not accelerate the negative age effect in the main regression model for the group comparison. Finally, there are a few putamen observations of PD patients that exceed the ones of healthy controls (**Figure S4**), that might have also lifted the PD intercept in putamen.

This finding requires additional validation because in PD, the D_2_R changes in putamen are dynamic and dependent on the disease stage. Compensatory D_2_R upregulation in putamen changes to downregulation around four years from PD onset [52, 54]. The PD duration greatly varied between the studies subjects (range: 0-29 years) and our findings cannot directly discern the effects in early and late stages, as the disease duration information was available for only half of the patients. Thus, we are unable to specify the weight of early and late stage patients in our data and analysis. For the subjects with available information, the correlation between disease duration and BP_ND_ was subtle but nevertheless negative in all tested regions (−0.17 in putamen) suggesting no prolonged upregulation with progressed PD. Consequently, our findings do not provide clear evidence against late stage D_2_R downregulation in putamen. Overall, the findings nonetheless highlight the region and motor symptom specificity of within-region D_2_R changes.

### Lowered between-region connectivity in severe violent behavior

Dopaminergic changes in severe violent behavior were more prominent in the regional coupling than in the overall regional receptor availability. Although in all groups we observed that thalamus showed the lowest regional connectivity, this hypoconnectivity was most salient in the subjects with severe violent behavior, who also had lowest regional coupling within the striatum. Aberrant coupling in this striatal dopamine network might reflect altered communication and weakened links between the regional nodes. Alternatively, the weakened correlation of D_2_R availability between different regions might also be explained by selective alteration of one specific dopaminergic region, while other regions are unaffected (without interaction between regions). However, this hypothesis was not notably supported by the regional analysis where the binding in subjects with severe violent behavior did not clearly differ from healthy controls in any of the regions.

We found some support for the similarity in the regional coupling patterns (correlation matrices) between the groups, except for group comparisons including severe violent behavior and pathological gambling. However, as the matrices we compared only had six individual data points (correlation estimates of 1) caudate-accumbens, 2) caudate-putamen, 3) caudate-thalamus, 4) accumbens-putamen, 5) accumbens-thalamus, 6) putamen-thalamus), the tests might be underpowered to show significant similarity between the matrices even if similarity exists, particularly if the reliability of the six input correlations (in the matrices) are calculated from a small sample size.

Altogether, we stress that the sample size for violent offenders was small which might limit the generalizability of our findings, although some of the other groups showing stronger correlation pattern were also small. Our findings nevertheless suggest different dopaminergic connectivity profiles between motor symptoms and violent aggression. The between-region connectivity in violence, and affective disorders such as schizophrenia, pathological gambling, and depression (for which we had limited sample sizes), should be further examined in larger studies.

### Unaltered baseline D_2_R in overweight

We observed neither regional D_2_R changes nor aberrant regional coupling in schizophrenia, pathological gambling, depression, or overweight groups. While some of the effects in the first three might have been overlooked due to limited sample sizes, overweight group included almost a hundred subjects. Our results, thus, underline that dopaminergic mechanisms do not notably contribute to general weight gain (without any subgrouping). Our findings accords with previous empirical studies [26, 55] as well as meta-analyses [56] finding no clear connections between altered D_2_R availability and obesity.

Finally, our results highlight that motor symptoms, rather than affective disorders or weight gain, are most consistently linked with regional alterations in D_2_R availability. According to our data, PD is associated with changes in regional D_2_R availability, while severe violent behavior might link to disturbed correlation between the regional availabilities. Overall, our results of unaltered D_2_R in overweight suggests that the postsynaptic type 2 dopamine receptor in the striatum and thalamus is not a central contributor of general weight gain, and possibly neither to schizophrenia, depression nor pathological gambling, although more data might be needed to clarify the issue. Overall, this large multi-group dataset suggests that the dopaminergic changes observed in neuropsychiatric pathology are multifaceted, involving region-specific and between-region mechanisms of the D_2_R. We conclude that the changes in baseline D_2_R might be more profoundly linked with motor symptoms and violent aggression than with affective disorders or weight gain.

### Effects of age and sex on D_2_R availability

We found that baseline striatal D_2_R availability is lowered through age both in healthy controls and in the groups of interest. Previously, we observed sex differences (female > male) in healthy controls consistently in the striatothalamic regions [26], whereas here in the groups of interest, sex differences were only found in the thalamus. However, this effect pertains to the joint analysis of the groups of interest together. Because all the groups did not include both males and females, we pooled the groups together for the analysis of age and sex, which might have overlooked group specific sex differences. Nevertheless, our data indicate that the effect of age on D_2_R is more consistent than the effect of sex, and generally found also in clinical populations. Accordingly, adjusting for age and sex is necessary when assessing the effects of other variables on striatothalamic D_2_R availability.

## Limitations

Our data were not fully balanced. Although the dataset was large, the group size, age, sex, and scanner distribution were compromised in some of the studied groups. Combined with our statistical analysis of the regional group differences that adjusted for the age, sex, and scanner related variation in the data, some group differences were possibly left undetected [51]. In the between-region analysis, our findings mainly indicate that in the healthy control, PD, and overweight groups, where we had sufficient data, the between-region binding correlation is strong. Larger samples are required to validate whether the striatothalamic correlation pattern is similarly strong or decoupled in the other groups with limited sample sizes. In a single PET scan, it is not possible to demonstrate the exact molecule-level mechanism for altered receptor availability. Not all subjects were drug naïve. Due to the retrospective nature of the data derived from multiple projects, the collection of clinical characteristics of patient populations were not standardized. Patient categorization was based on their enrollment criteria in the original studies. Finally, it must be noted that unaltered baseline D_2_R level does not mean the dopaminergic responses in different environmental contexts would be similar across all studied groups. It is possible that differential patterns of dopamine firing in motivationally triggering environments exist for example, towards gambling in pathological gamblers and towards feeding in obese individuals [25]. Investigating such effects however requires standardized challenge or activation paradigms, which could not be incorporated in the current analysis.

## Conclusions

Dopaminergic function is altered across multiple psychiatric and neurological conditions. The regional receptor availability patterns, as well as interregional coupling of D_2_R levels, might distinguish between the specific conditions. PD was associated with region-specific changes in D_2_R availability. The role of D_2_R in violent behavior might rather relate to the interregional connections, as in this subject group, the between-region receptor availabilities were less correlated. In pathological gambling and overweight groups, no clear changes were observed in the striatothalamic D_2_R level nor the correlation structure. Our results suggest that in striatum and thalamus, motor symptoms are more profoundly linked to the region-specific D_2_R availability, while violence might be more associated with lowered correlation between the regions.

## Acknowledgements

The study was supported by the Päivikki and Sakari Sohlberg Foundation (grants to T.M. and V.K.), the State research funding for expert responsibility area (ERVA) of the TYKS Turku University Hospital (grants to T.M. and J.T.), the Sigrid Juselius Foundation (grants to L.N. and J.R.), and Academy of Finland (grant numbers 294897 and 332225 to L.N., and grant number 310962 to J.R.), Finnish Governmental Research Funding (VTR) for Turku University Hospital (grant to J.R.), and the Finnish Cultural Foundation (grant to V.K.). We acknowledge Abbvie, Nordic Infucare; Advisory Board: Abbvie, Nordic Infucare, Adamant Health Ltd (Honoraria for Lecturing to V.K.). We thank Jose Manuel Rivera Espejo and Joni Virta for sharing their expertise on statistical modeling regarding this study. We also acknowledge Tomi Karjalainen for his work in image processing and statistical modeling, Tuomas Knuuti for quality control, and Veera Korhonen for literature searches.

## CRediT authorship contribution statement

**Malén**: Conceptualization, Methodology, Validation, Formal analysis, Data curation, Writing -Original Draft, Writing – original draft, Writing – review & editing, Visualization, Project administration.

**Santavirta**: Conceptualization, Methodology, Writing – review & editing, Visualization.

**De Maeyer**: Methodology, Software, Writing – review & editing, Visualization.

**Tuisku**: Methodology, Software, Validation, Formal analysis, Data curation, Writing – review & editing, Visualization.

**Kaasinen**: Conceptualization, Investigation, Resources, Data Curation, Writing – review & editing.

**Kankare**: Investigation, Resources, Data Curation, Writing: Review & Editing.

**Isojärvi**: Software, Data curation, Writing – review & editing.

**Rinne**: Investigation, Resources, Writing – review & editing.

**Hietala**: Investigation, Resources, Writing – review & editing.

**Nuutila**: Investigation, Resources, Writing – review & editing.

**Nummenmaa**: Conceptualization, Investigation, Resources, Writing – review & editing, Visualization, Supervision, Funding acquisition.

## Data availability

The data were retrieved from the Turku PET Centre Aivo database (https://aivo.utu.fi). We present the mean brain maps of group specific D_2_R availabilities in NeuroVault (https://neurovault.org; https://identifiers.org/neurovault.collection:12799).

## Ethics approval

Finnish legislation does not require ethical approval for register-based studies.

