## Supplementary material for "Alterations in type 2 dopamine receptors across neuropsychiatric conditions: A large-scale PET cohort"

**Original publications whose data are used in the current study** (not comprehensive)

Hietala J, Syvälahti E, Vuorio K, Någren K, Lehikoinen P, Ruotsalainen U, Räkköläinen V, Lehtinen V, Wegelius U. Striatal dopamine D2 receptor characteristics in neuroleptic-naive schizophrenic patients studied with positron emission tomography. Arch. Gen. Psychiatry 51: 116-123, 1994.

Tiihonen J, Kuoppamäki M, Någren K, Bergman J, Eronen E, Syvälahti E, Hietala J. Serotonergic modulation of D2 dopamine receptor binding in healthy volunteers in vivo, Psychopharmacology, 126: 277-280, 1996.

Hietala J, Någren K, Lehikoinen P, Ruotsalainen U, Syvälahti E. Measurement of striatal D2 dopamine receptor density and afinity with 11C-raclopride - a test-retest analysis. J Cereb Blood Flow Metab 19:219-217, 1999.

Hagelberg N, Någren K, Kajander J, Hintikka S, Hietala J, Scheinin H. μ-opiate receptor agonism increases dopamine D2 receptor binding in human basal ganglia – a positron emission tomography study. Synapse 45(1):25-30, 2002

Aalto S, Hirvonen J, Kajander J, Scheinin H, Någren K, Vilkman H, Gustafsson L, Syvälahti E, Hietala J. Ketamine does not decrease striatal dopamine D2 receptor binding in man. Psychopharmacology (Berl) 164(4):401-406, 2002.

Hirvonen J, Aalto S, Lumme V, Någren K, Kajander J, Vilkman H, Hagelberg N, Oikonen V, Hietala J. Measurement of striatal and thalamic dopamine D2 receptor binding with [11C]Raclopride, Nucl Med Comm, 24:1207-1214, 2003.

Penttilä J, Kajander J, Aalto S, Hirvonen J, Någren K, Ilonen T, Syvälahti E, Hietala J. Effects of fluoxetine on dopamine D2 receptors in the human brain: a positron emission tomography study with 11Craclopride. Int J Neuropsychopharmacol 7: 431-439, 2004.

Hirvonen J, van Erp TGM, Huttunen J, Någren K, Aalto S, Huttunen M, Lönnqvist J, Kaprio J, Hietala J, Cannon TD. Increased Caudate Dopamine D2 Receptor Availability as a Genetic Marker for Schizophrenia. Arch Gen Psychiatry, 62(4):371-8, 2005.

Hirvonen J, Karlsson H, Kajander J, Markkula J, Rasi-Hakala H, Någren K, Salminen JK, Hietala J. Striatal dopamine D2 receptors in medication-naive patients with major depressive disorder as assessed with [11C]raclopride PET. Psychopharmacology (Berl). 2008 May;197(4):581-90.

Hietala, J, Mika Hirvonen, , Jukka Huttunen, Harry Vilkman, Jussi Hirvonen, , Ville Lumme, , Kjell Någren, Pertti Neuvonen, Mikko Niemi, Raimo K.R. Salokangas. Optimization Of Antipsychotic Drug Treatment With Perphenzine –A Pharmacogenetic And Positron Emission Tomography Study. Submitted.

| | N | GE Advance | HR + | HRRT | GE* VCT PET/ CT | GE* 690 PET/CT | Ecat 931 |
|---|---|---|---|---|---|---|---|
| Healthy controls | 239 | 69 | 13 | 81 | 9 | 17 | 50 |
| Parkinson's disease | 60 | 0 | 6 | 0 | 0 | 0 | 54 |
| Schizophrenia | 7 | 3 | 0 | 0 | 0 | 0 | 4 |
| Violence | 10 | 0 | 0 | 0 | 10 | 0 | 0 |
| Pathological gambling | 12 | 12 | 0 | 0 | 0 | 0 | 0 |
| Depression | 12 | 12 | 0 | 0 | 0 | 0 | 0 |
| Overweight | 97 | 38 | 6 | 16 | 8 | 27 | 2 |

**Table S1**. The number of subjects for each scanner in the sample. N= Number of subjects in total. GE*= GE Discovery.

#### $D_2R$ availability through PD duration

To assess how $D_2R$ availability changes through PD duration, we calculated their correlation for each region in the subset of PD patients who had the disease duration information available (n= 27). Shapiro-Wilk test supported the normal distribution of the log-transformed binding potential estimates (p-value > 0.2 in each region). Thus, we used the log-transformed binding and Pearson correlation in the analysis. The test was conducted as one-tailed, because we expected negative correlation if any (binding decreases as disease progresses), due to the degenerative nature of the disease. The analyses of normality and correlation were run in R with the package stats [1]. The correlation between PD duration and log-transformed binding was negative in all four regions: -0.44 in caudate (p= 0.01), -0.28 in accumbens (p= 0.08), -0.17 in putamen (p= 0.2), and -0.34 (p= 0.04) in thalamus. Although age was not adjusted for in this analysis, the finding suggest that the $D_2R$ availability decreases as PD progresses (**Figure S1**).

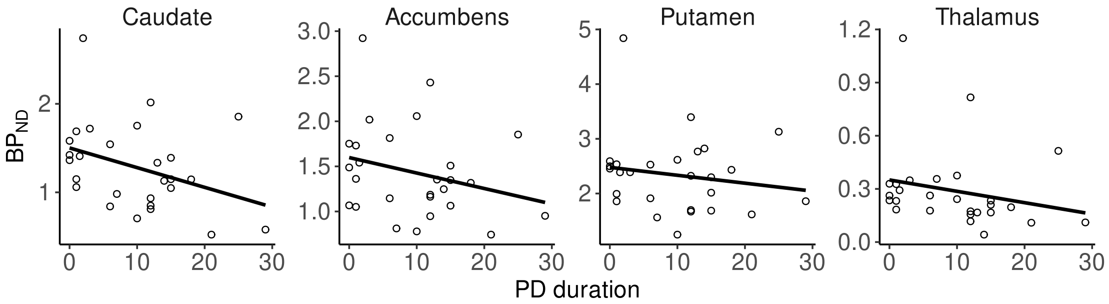


**Figure S1**. The $BP_{ND}$ estimates throughout the PD duration in years (mean 10 and median 9 years, range from 0 to 29 years, for those subjects who we had the information, n=27).

| | N | Mean | SD | Range | NA |
|---|---|---|---|---|---|
| Healthy controls | 147 | 22.3 | 1.6 | 17.7-25.0 | 92 |
| Parkinson's disease | 11 | 26.8 | 5.1 | 20.3-39.4 | 49 |
| Schizophrenia | 2 | 26.9 | 3.0 | 24.1-29.7 | 5 |
| Violence | 10 | 27.2 | 2.3 | 24.0-30.7 | 0 |
| Pathological gambling | 12 | 27.8 | 3.4 | 21.2-35.9 | 0 |
| Depression | 12 | 26.5 | 5.2 | 18.9-36.5 | 0 |
| Overweight | 97 | 31.5 | 6.5 | 25.1-49.0 | 0 |

**Table S2**. Body mass index for the 291 subjects (67% of the total sample) for who we had the information available. N= number of observations. NA= Number of subjects with missing BMI information.

### Validation of PET atlas based spatial normalization of the [$^{11}$C]raclopride binding estimates

Two alternative spatial normalization methods can be used to calculate binding potential ($BP_{ND}$) estimates from a [$^{11}$C]raclopride PET image: i) a deformation-field based method with Freesurfer (https://surfer.nmr.mgh.harvard.edu/) using the subject's magnetic resonance image (MRI) [2], and ii) PET atlas template-based normalization with SPM (https://www.fil.ion.ucl.ac.uk/spm/) and an in-house atlas. As for the whole sample we did not have MRI available, we used the PET atlas method to maximize sample size. For validation, however, we compared the two methods. Out of the total sample (n=437), 249 subjects had MRI available. For these subjects, we assessed the similarity of their alternative $BP_{ND}$ estimates.

To assess, how similar estimates the two normalization methods produce, we calculated the correlation between their original-scale binding estimates. We first tested the normality of the binding estimates, separately for each region within both normalization methods. As the Shapiro-Wilk test did not support normality of the binding estimates in all the tests, we used Spearman correlation. Spearman correlation was applied as one-tailed, because the variables to compare are measures of binding, and thus we expected to test only positive correlation, which was also supported in our previous work in healthy controls [3]. The analyses were run in R with the package stats [1].

As expected, the correlation of the alternative $BP_{ND}$ estimates was strongest in the striatum where the reliability of [$^{11}$C]raclopride binding is the highest [e.g. 4], and despite a few outlier observations, the methods produce consistent estimates also in thalamus (**Figure S2**). The correlation was 0.88 ($p< 0.001$) in caudate, 0.89 ($p< 0.001$) in accumbens, 0.92 ($p< 0.001$) in putamen, and 0.78 ($p< 0.001$) in thalamus. The average and difference of the two methods are presented in **Figure S3**. Overall, the MRI based estimates are higher than the PET atlas based estimates, except for accumbens where the observations are more evenly represented on both sides of the zero-difference.

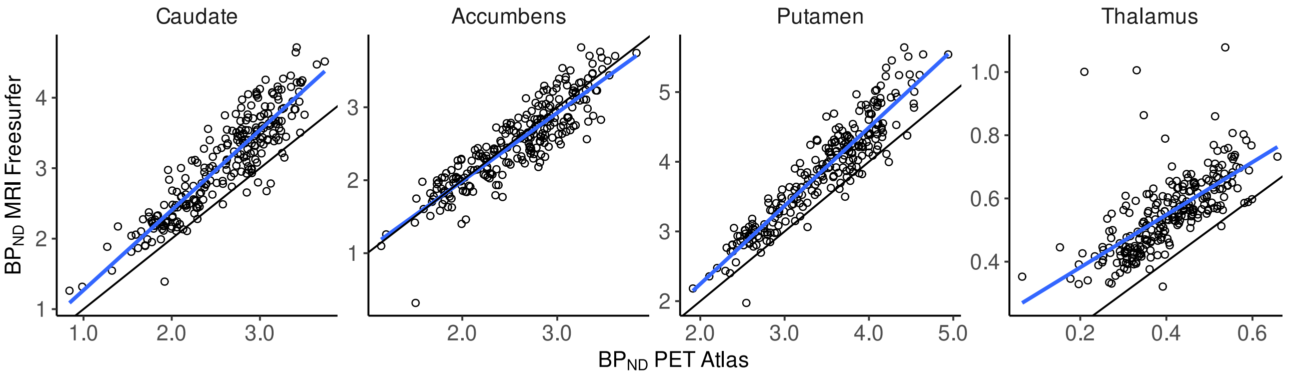


**Figure S2**. The association of regional [$^{11}$C]raclopride binding estimates from SPM with PET atlas template (x-axis), and Freesurfer with subject MRI (y-axis). The figure shows the observations of a subsample of 249 subject for who both methods could be applied. The lines represent the linear model (blue) and the y=x line (black).

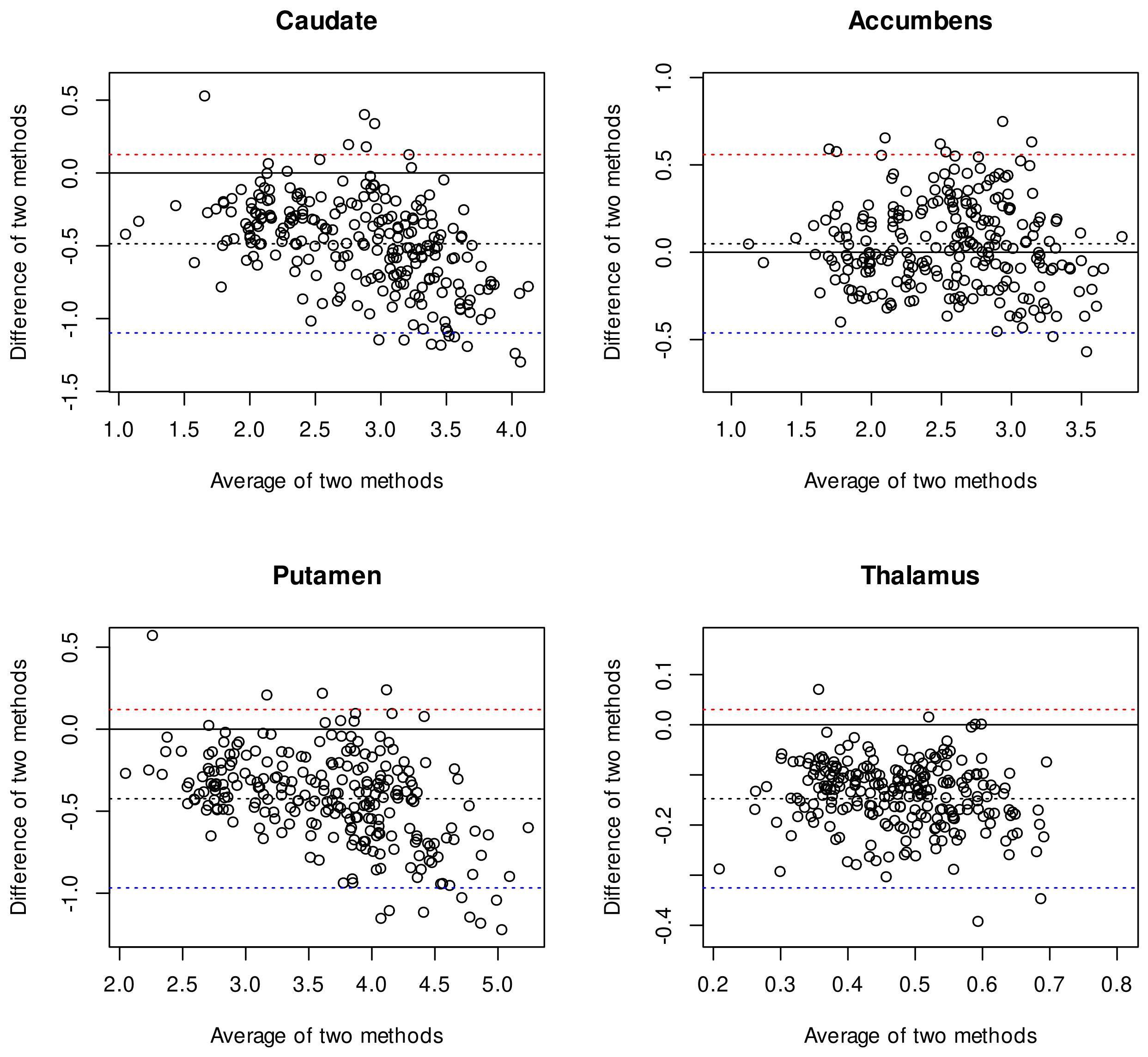


**Figure S3**. The association of original-scale regional [$^{11}$C]raclopride binding estimates from SPM with PET atlas template, and Freesurfer with subject MRI. The Bland Altman plot [5] shows the average and the difference (PET atlas – MRI based estimate) of the two methods. The figure shows observations of a subsample of 249 subject for who both methods could be applied. The solid line (black) indicates zero-difference. The dashed lines represent the mean (black), the upper limit (red), and the lower limit (blue). The limits= Mean of difference + or - (1.96 × standard deviation of difference).

| | Caudate | | Accumbens | | Putamen | | Thalamus | |
|---|---|---|---|---|---|---|---|---|
| | Mean ratio | Diff range | Mean ratio | Diff range | Mean ratio | Diff range | Mean ratio | Diff range |
| Healthy controls | 1.02 | -0.40, 0.54 | 1.00 | -0.54, 0.92 | 1.02 | -0.21, 0.46 | 0.92 | -0.15, 0.09 |
| Parkinson | 1.14 | -0.36, 0.44 | 1.01 | -0.83, 0.64 | 1.01 | -0.59, 0.51 | 0.90 | -0.15, 0.09 |
| Schizophrenia | 0.97 | -0.32, 0.16 | 1.00 | -0.39, 0.16 | 1.05 | 0.01, 0.26 | 0.89 | -0.19, -0.00 |
| Violence | 1.00 | -0.25, 0.15 | 1.00 | -0.38, 0.39 | 1.01 | -0.10, 0.15 | 0.89 | -0.08, 0.01 |
| Gambling | 0.99 | -0.14, 0.13 | 0.94 | -0.29, 0.02 | 1.00 | -0.17, 0.12 | 0.85 | -0.12, -0.01 |
| Depression | 1.03 | -0.01, 0.19 | 1.04 | -0.14, 0.31 | 1.02 | -0.06, 0.11 | 0.90 | -0.12, 0.04 |
| Overweight | 1.01 | -0.20, 0.27 | 0.98 | -0.71, 0.48 | 1.01 | -0.16, 0.59 | 0.93 | -0.13, 0.06 |

**Table S3**. Group-specific assessment of hemispheric symmetry. Mean ratio= mean of within-subject original-scale $BP_{ND}$ ratios (left / right hemisphere). Diff range= range of the absolute within-subject original-scale $BP_{ND}$ differences (left – right hemisphere).

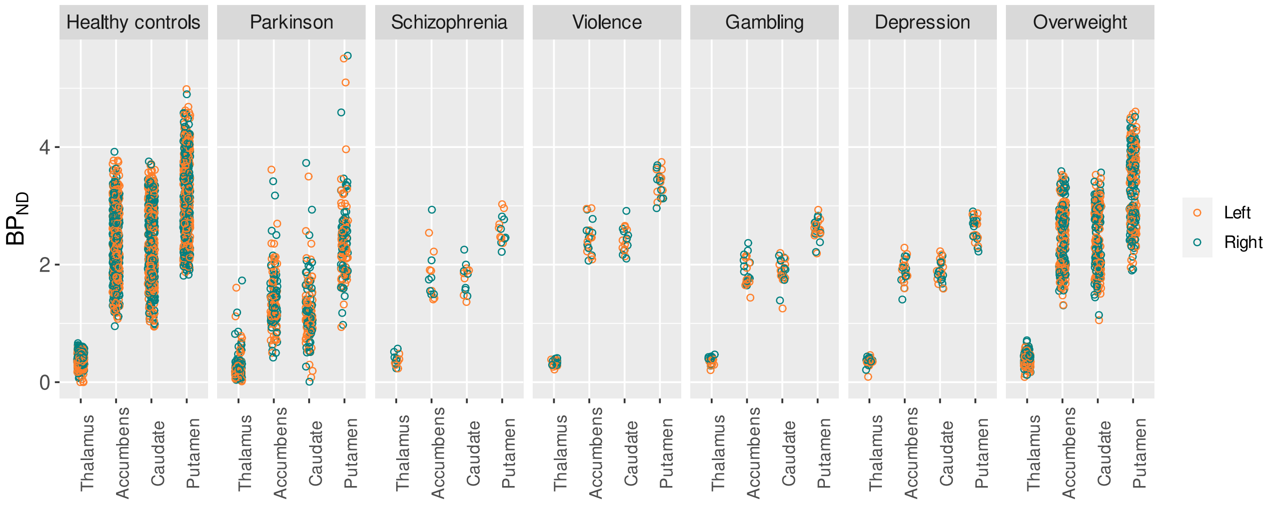


**Figure S4**. Left and right original-scale $BP_{ND}$ estimates separately for each group.

### Model diagnostics and comparison

In the main model, ROI was modeled as a random intercept and as a random slope for group, age, and sex effects (**Model 1**). When ROI is modeled as a random (and not fixed) effect, the model can utilize the information of the other ROIs to improve the estimates of each ROI and the non-independence of the observations across ROIs [6]. As a result, the regionally varying effects are partially pooled across ROIs (shrinkage towards the mean) [6-8], which is also considered as a way to correct for multiple comparisons [9]. However, we only have four ROIs, while >5 levels of the random variable are recommended, and the ROIs are distinguishable units that constitute the whole population of interest (compare a random sample from a population of interest) [6]. As a result, modeling ROI as a fixed effect would also be justified [6]. Hence, we ran additional models (**Models 2-4**) with ROI as a fixed effect. More precisely, to get ROI-specific group effects, ROI was modeled as an interaction with group (instead of as a single fixed effect) in the alternative models.

Model 1 (main model):

log_bp ~ 1 + group + age_z + male + (1 | subject) + (1 | gr(scanner, id = "scanner")) + (1 + group + age_z + male | gr(roi, id = "roi")) + (1 | gr(scanner:roi, id = "scanner:roi")),

sigma ~ (1 | gr(scanner, id = "scanner")) + (1 | gr(roi, id = "roi")) + (1 | gr(scanner:roi, id = "scanner:roi"))

Model 2:

log_bp ~ 1 + age_z + male + group*roi + (1 | subject) + (1 | gr(scanner, id = "scanner")),

sigma ~ (1 | gr(scanner, id = "scanner"))

Model 3:

log_bp ~ 1 + age_z + male + group*roi + (1 | subject) + (1 | gr(scanner, id = "scanner")) + (1 | gr(scanner:roi, id = "scanner:roi")),

sigma ~ (1 | gr(scanner, id = "scanner")) + (1 | gr(scanner:roi, id = "scanner:roi"))

Model 4:

log_bp ~ 1 + age_z + male + group*roi + (1 | subject) + (1 | gr(scanner:roi, id = "scanner:roi")),

sigma ~ (1 | gr(scanner:roi, id = "scanner:roi"))

**Models 1-4**. Basic structure of the models using syntax of brms [10-12] utilizing rstan [13] in R [1]. To help modeling convergence, maximum treedepth was set to 20 and adapt delta to the range of 0.99-0.999 (https://mc-stan.org/misc/warnings.html#divergent-transitions-after-warmup). Male indicates sex (male, female). Age_z refers to standardized age, and subject to subject index. Syntax 'gr' refers to basic grouping structure (https://rdrr.io/cran/brms/man/gr.html), applied explicitly to random variables (roi, scanner, roi:scanner) also applied to residual variances (sigma).

*Main model diagnostics.* Supporting sufficient convergence of the main model, there were no divergent transitions and the Rhats were 1 [10]. To produce generalizable results and to avoid overfitting, we estimated how well the main model predicts new observations by running k-fold (k=3) cross-validation using brms and loo [10-14] packages in R [1]. The cross-validation supported good specification of the main model, as the total number of parameters (here 609) was larger than the estimated effective number of parameters (p_kfold, here 534, standard error 38.3) [10-12, 15] indicating that the results generalize sufficiently well to unobserved data.

*Model comparison.* Additionally, we compared the models with the 3-fold and pareto smoothed importance sampling (PSIS) leave-one-out cross-validation [16] with loo package [14]. Based on the cross-validation estimates (elpd_diff and se_diff) described by Vehtari and colleagues [16], the poorest fit was Model 2, where we excluded the random effect of combined scanner and ROI (scanner:roi with 6 scanners x 4 ROIs = 24 levels). Overall, the cross-validation did not show clear signs of the alternative models being stronger at predicting new data. Thus, we reported the results of the main model specifying ROI as a random factor, coherently with our previous work [3].

**Validation of the age and sex effects on regional $D_2R$ availability**

To test whether the observed effects of age (negative) and sex (higher binding in females than males) generalize outside the healthy control sample [3], we calculated these effects in two separate models predicting the $BP_{ND}$ in i) healthy controls (n= 239, 174 males, 65 females, almost identical to the sample in our earlier work [3]), and ii) the groups of interest, including the subjects with Parkinson's disease, schizophrenia, pathological gambling, depression, past severe violent behavior, and overweight (n=198, 120 males, 78 females). For these models, we included only age and sex (and not group) as fixed effects. However, in the model analyzing the groups of interest, we allowed the intercept to vary by group (random intercept for group) to avoid the model from interpreting group-related variance in $BP_{ND}$ as an age or sex effect. The $D_2R$ availability decreased through age both in healthy controls about 10% in 14 years, and in the clinical groups about 10% in 17 years (**Figure S5**). On average, the decrease was about 10% in one SD ≈ 15 years (SD was 14 years in healthy controls and 17 years in the clinical groups). The age range was 19-82 in both healthy controls and in the clinical groups. Most of the probability mass supported higher binding in females than males in the healthy controls, but the difference between the sexes in the striatum was not clear in the clinical data. In thalamus, however, all subjects showed support for higher binding in females than males. Females had approximately 5% (healthy controls) and 10% (clinical subjects) higher thalamic binding than males.

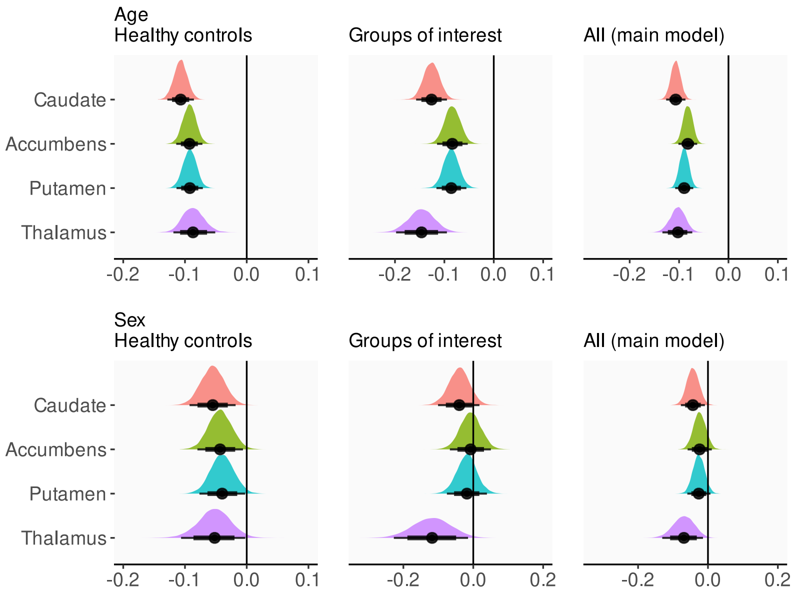


**Figure S5**. The effects of age (upper row) and sex (bottom row) separately among the healthy controls, the other subjects (groups of interest), and all subjects (from the main model). The x-axis shows regression coefficient of the log-transformed $BP_{ND}$. The effect of sex is presented as male - female, meaning negative values support higher $BP_{ND}$ of females versus males.

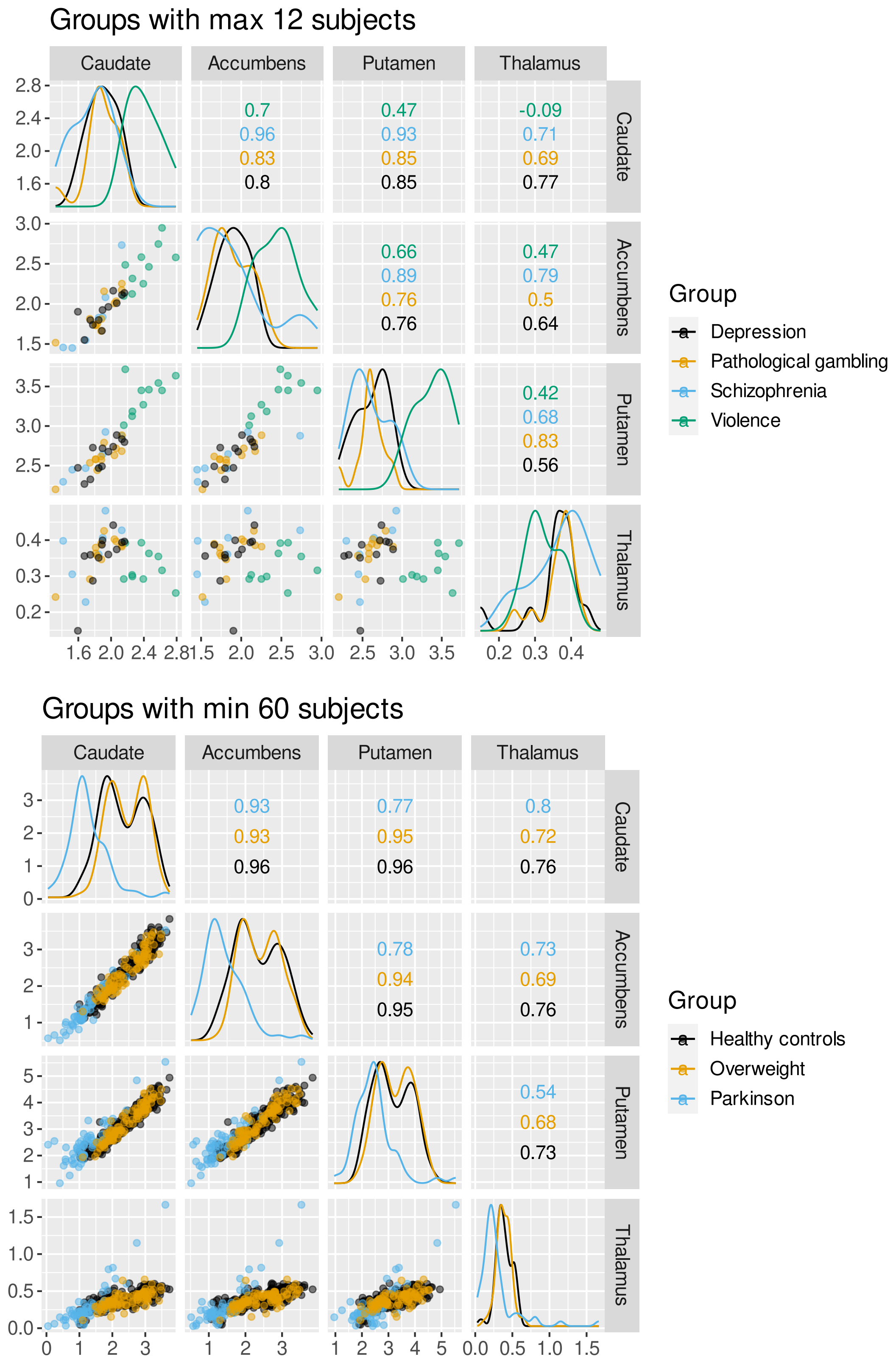


**Figure S6.** Between-region scatter plots (lower triangle) and correlations (upper triangle), as well as regional density plots (diagonal axis) of the original-scale $BP_{ND}$ estimates, separately for each subject group. Small groups (maximum of 12 subjects) and larger groups (at least 60 subjects) are presented separately to enhance the visibility of the observations and their relationship between x and y axes.
